# An Online Brain-Computer Interface for Detecting Incongruity in Augmented Reality Applications

**DOI:** 10.64898/2025.12.20.694925

**Authors:** Michael Wimmer, N. ElSayed, Bruce H. Thomas, Gernot R. Müller-Putz, Eduardo E. Veas

**Affiliations:** Know Center Research GmbH, Graz, Austria; Institute of Neural Engineering, Graz University of Technology, Graz, Austria; Wearable Computer Lab, University of South Australia, Mawson Lakes, SA, Australia; BioTechMed Graz, Graz, Austria; Institute of Human-Centred Computing, Graz University of Technology, Graz, Austria

**Author notes:** Author to whom any correspondence should be addressed.

**Keywords:** Electroencephalography (EEG), Augmented reality (AR), Event-related potential (ERP), N400, Eye tracking

## Abstract

**Objective:** Augmented reality can provide digital information about physical entities presented within its real-world context. However, this information might disagree with the user’s expectations due to factual errors in the data or cognitive biases. Such incongruity can impair user experience and undermine trust in the AR system. To address this issue, we propose detecting inconsistencies between physical objects and digital information through hybrid brain-computer interfaces.

**Approach:** We conducted two complementary experiments. First, we implemented a strategy that integrates eye-tracking and brain signals for incongruity detection in an offline study. Subsequently, we assessed our approach in an online study in which participants received immediate feedback on the classification.

**Main results:** The grand average event-related potentials revealed consistent electroencephalographic responses to incongruent augmentations, specifically a centroparietal N400 effect, across both experiments. We could further distinguish between congruent and incongruent information with an average balanced accuracy of 70 % in the online study.

**Significance:** These findings demonstrate the feasibility of detecting incongruity online, allowing for autonomous system adaptation, like presenting information in a more accessible format or providing contextual support.

## 1 Introduction

Augmented reality (AR) allows users to simultaneously experience and interact with the physical and virtual worlds in real time using [1]. Situated visualizations present virtual information directly within their real-world context, e.g., using see-through head-mounted displays (HMDs). This fosters an intuitive connection between a user’s physical environment and digital data [2]. Locating data in the spatial proximity of physical entities communicates additional information to users and raises the opportunity for in-situ sensemaking [3, 4]. However, such in-situ presented information does not necessarily agree with the expectations of the user. Discrepancies could arise from factual errors in the underlying data, for instance, when drawing from outdated or unreliable sources. On the other hand, information that contradicts a user’s prior knowledge or experience, or is misinterpreted, may result in perceived errors despite being accurate. Such incongruity between the physical world and related virtual data can negatively impact the user experience and trust in the technology [5]. To address these effects, we propose to autonomously detect incongruent AR information online (i.e., in real-time) based on users’ physiological data through a hybrid brain-computer interface (BCI) [6, 7].

Previous research has identified neurophysiological correlates associated with the processing of incongruent information using electroencephalographic (EEG) signals. Specifically, the N400 is a component of event-related potentials (ERPs) characterized by a negative-going deflection that occurs approximately 200 to 600 ms after the presentation of meaningful, incongruent stimuli [8]. The concept of incongruity is specific to each person’s perspective and current mental state, yet the N400 has been observed across diverse contexts. Initially, this component was linked to language processing. In a sentence-reading task, words were presented one at a time, with the final word occasionally being out of context (“He took a sip from the *transmitter* ”) [9]. Subsequently, the N400 effect was observed for a variety of stimulus types and modalities, including visuals [10], sounds [11], and mathematical equations (“7 x 4 = *26* ”) [12].

Recently, incongruity was also studied within the context of AR [13]. This work characterized the neurophysiological responses to processing mismatches between a physical referent and related AR information. In a controlled experiment, the information was presented in a predefined and cued location known to participants. This eliminates eye movements affecting the brain signals and enables straightforward computation of incongruity-related EEG responses. The study revealed that (i) the N400 effect is also modulated by AR mismatches and that (ii) offline synchronous (i.e., cue-based) decoding is feasible above chance (approximately 65 % accuracy). However, the implemented BCI approach is considerably limited in its practical usability: In real-world AR applications, digital elements are not presented at cued positions but are organically embedded in the physical environment [4, 14]. AR users freely deploy their gaze across the environment to find information that is relevant to them. Hence, the synchronous classification strategy (time-locked to the presentation of the information) presented in [13] is not transferable to practical AR use cases with online classification.

Similarly, another work investigated contextual expectations by presenting AR objects in incongruent physical environments [15]. While this study demonstrated the feasibility of observing consistent incongruity-related brain responses also outside the lab, it suffered from the same practical limitations in user and gaze behavior and AR object integration as [13].

Online BCIs designed for practical applications [16] need to decode brain signals without external cues (as they rarely exist in out-of-the-lab scenarios, if at all). Therefore, asynchronous BCIs [17, 18] continuously analyze and classify the ongoing EEG signals and discriminate brain patterns of interest from noise and task-irrelevant stimuli. This naturally increases the number of false positive detections compared to synchronous BCIs, which restricts the analysis to fixed windows after stimulus presentation [19, 20]. Consequently, decoding the N400 effect in an online scenario requires a strategy to minimize false detections. To address this, our approach is to incorporate natural gaze behavior monitored by the eye tracker integrated into the HMD. Using eye tracking, the system can detect when users shift their gaze toward an AR visualization, providing a real-time marker to align the EEG responses. In this way, brain signals are linked to the currently viewed content. Rather than continuously decoding the ongoing brain activity, this strategy restricts the analysis to a fixed time window triggered by the user’s gaze at an augmentation. This sequential procedure (eye tracker informs BCI) allows us to decode incongruity without restricting users’ natural gaze behavior. To the best of our knowledge, this is the first demonstration of online decoding of incongruent information using any presentation modality.

Successfully detecting incongruity online contributes to the development of autonomously adaptive AR systems [21], which could assist users by proactively providing additional contextual information and explanations or by presenting information in a more accessible manner. Beyond individual assistance, online detection could help diagnose systems by automatically flagging incongruities, facilitating robust system designs without requiring explicit user feedback [22].

This work addresses the research gap between offline studies of EEG responses to incongruity in gaze-fixation tasks and the development of online N400-BCIs. Specifically, we investigated the following research questions (RQs):

**(RQ1)** Can eye tracking be used as a marker to reveal the EEG responses to incongruent AR information, and can they be decoded via machine learning models?
**(RQ2)** Can AR incongruity be detected online using EEG-based machine learning models?

To develop an online N400-BCI for AR applications, we conducted two experiments. In *Experiment 1*, participants performed an interactive task reflecting a typical use case of AR, i.e., providing additional information about physical objects through situated visualizations. Congruent and incongruent numerical data were presented in the referent’s proximity, requiring participants to actively shift their gaze to process the information. In a hybrid BCI approach, we informed EEG-based classifiers with the eye tracker to perform offline incongruity decoding. In *Experiment 2*, we applied the findings from *Experiment 1* and detected incongruities online. Participants received visual feedback based on the classification output.

## 2 Experiment 1

### 2.1 Methods

#### 2.1.1 Participants

Fifteen participants were recruited for this experiment (28.8 ± 2.9 years (mean (M) ± standard deviation (SD)), 10 male, 5 female). All participants were free of known neurophysiological disorders and had normal or corrected-to-normal vision. Additionally, all participants completed an online Ishihara pseudoisochromatic test^1^ to assess their color vision. Two participants (13 %) reported having no prior AR experience, 8 reported being beginners or intermediates (53 %), and 5 reported being advanced or experts (33 %). Further, a total of 11 participants (73 %) had previously participated in at least one EEG study. This experiment was conducted in accordance with the Declaration of Helsinki (1975) and approved by the ethics committee of Graz University of Technology. All participants were provided with written and oral instructions regarding the purpose and procedure of the study and signed an informed consent form. Participants received vouchers worth 20 euros as compensation.

#### 2.1.2 Experimental Setup and Procedure

Participants were seated at a table such that they could comfortably reach and manipulate a tricolor (blue, white, and red) Rubik’s Cube (Figure 1b and Figure 2). Using a platform, the cube’s top face was oriented towards the participant at a 30° angle to the table. A camera (Renkforce RF AC4K 300, Conrad Electronic SE, Hirschau, Germany) was aimed at the cube to capture the nine colored squares that made up its top face. The implemented color detection algorithm was inspired by the Qbr Rubik’s Cube solver^2^. A HoloLens 2 HMD (Microsoft, Redmond, WA, USA) was utilized to present the AR visualizations created in Unity 2021.1.31.

The experimental procedure lasted approximately 1.5 hours and included three blocks. In the first block, participants familiarized themselves with the HMD and the experimental paradigm. The second and third blocks began with the calibration of the HMD’s eye tracking system and were divided by a break of about 15 minutes, during which participants could remove the headset. Both blocks consisted of 5 runs of 21 trials each. In each run of five to seven minutes, 14 congruent (67 %) and 7 incongruent (33 %) counts were presented in a pseudo-randomized order, resulting in a total of 210 trials (10 runs × 21 trials) per participant. Before the first block, we mounted the EEG electrodes, as described in Section 2.1.3.

**Figure 1.**
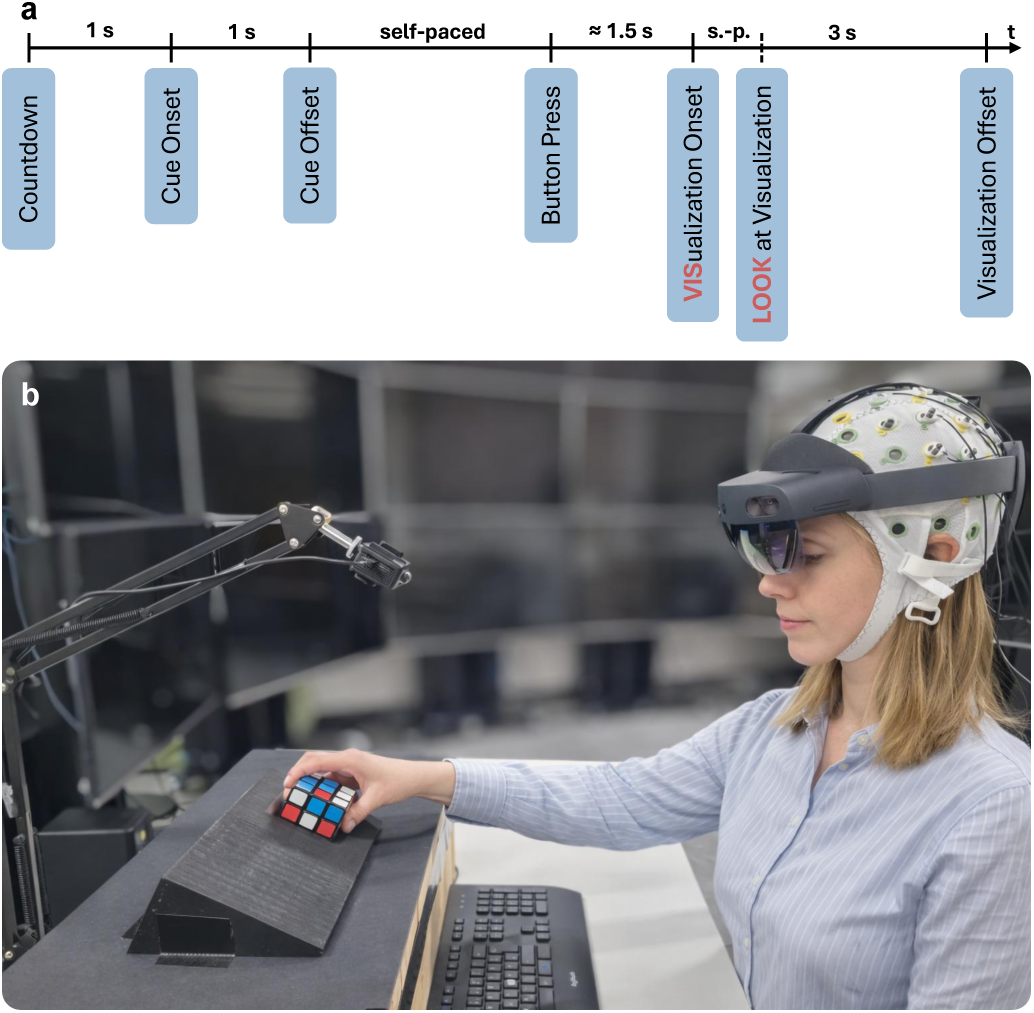
Experimental design of *Experiment 1*. **(a)** Timing of a single trial, including highlighted VIS and LOOK alignments. **(b)** A participant equipped with the EEG system and the HMD interacts with the Rubik’s Cube (image digitally recreated for privacy reasons).

Each trial commenced with a countdown from two to zero, signaling participants to prepare for the upcoming visual cue. A green arrow was displayed for a duration of 1 s, indicating the row or column to be rotated and the direction of rotation. Following the rotation, participants returned the cube to its initial position and counted the number of blue, white, and red squares on the top face. Once the correct count was determined, participants were instructed to press a physical keyboard button while focusing on the center of the cube. After a randomized delay of 1.25 to 1.75 s, this action triggered the display of a visualization showing either a **congruent** or an **incongruent** count in the order blue-white-red. Incongruent counts were inherently impossible, with one of the three digits differing by exactly three (e.g., 5-3-4 or 2-3-1 instead of 2-3-4). The range of each digit was fixed between 0 and 9. When participants realized the visualization appeared, they shifted their gaze from the cube to the displayed information. The visualization extended from 6.5 to 12 cm left of the cube, equivalent to an eccentricity range of 7.4 to 13.5° at a viewing distance of approximately 50 cm. They were free to naturally move their eyes across the scene (e.g., the visualization and the cube) until the visualization disappeared after 3 s. The subsequent trial began after a break of 1 s. To keep the overall duration of the experiment at a minimum, participants were not asked to indicate detected incongruities after the trials. A video can be found in the supplementary materials.

Participant 5 completed only 198 of 210 trials due to technical problems. The paradigm is adapted with permission from a previous study by Wimmer and colleagues [13]. Their paradigm can be accessed online via the link in the original paper.

#### 2.1.3 Data Acquisition

We measured EEG signals using 28 active electrodes (actiCAP, Brain Products GmbH, Glitching, Germany), positioned according to the international 10-10 systems at Fz, FC3, FC1, FCz, FC2, FC4, C5, C3, C1, Cz, C2, C4, C6, CP5, CP3, CP1, CPz, CP2, CP4, CP6, P3, P1, Pz, P2, P4, PO3, POz, and PO4. The ground and reference electrodes were placed at the left and right mastoids, respectively. This electrode layout covers frontocentral and centroparietal areas of the brain, which are particularly relevant for incongruity processing [8]. Four additional active electrodes were used to record electrooculographic (EOG) signals. Specifically, we positioned two electrodes at the superior and inferior canthi of the left eye, and two at the outer canthi of both eyes. EEG and EOG signals were recorded at a sampling frequency of 500 Hz. To ensure high signal quality, we kept the impedance between electrodes and scalp below 15 kΩ and continuously monitored the signals throughout the experiment. Additionally, we used the HMD’s eye tracker to determine, for each trial, at which time point the participants’ gaze first intersected with the AR visualization. Eye tracking data was sampled at 30 Hz. All data streams were recorded and synchronized utilizing the lab streaming layer protocol [23].

**Figure 2.**
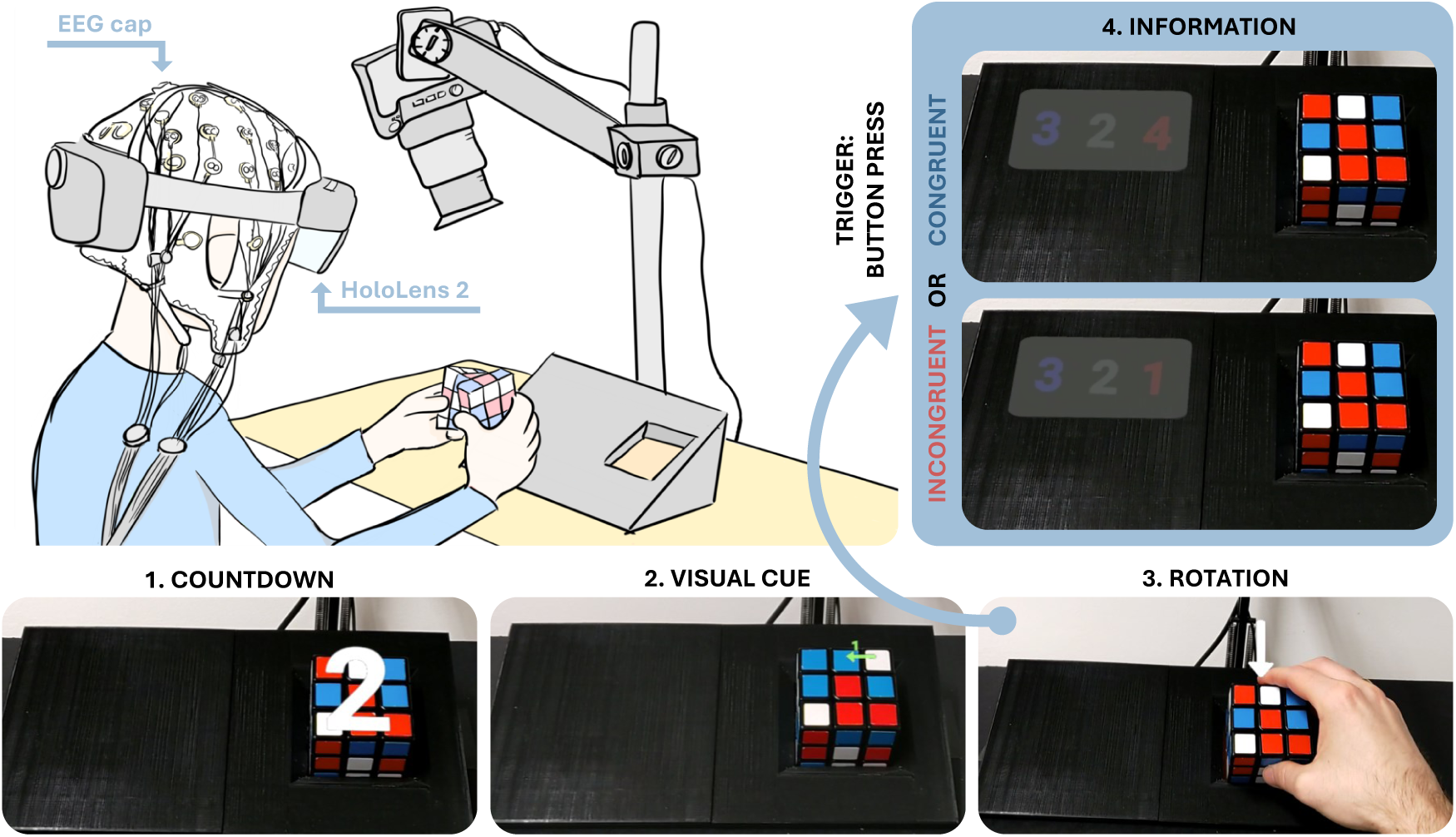
Experimental setup and design. **Top left:** Participant wearing an EEG cap and a HoloLens 2 while rotating the tricolor Rubik’s Cube. **(1)** Each trial begins with a countdown. The cube is in its starting position. **(2)** A visual cue indicates which row or column should be rotated. **(3)** Participants perform the rotation, return the cube to its starting position, and count the colors on the top face. **(4)** The information regarding the cube’s current status is triggered by a button press. In the example shown, the congruent answer is 3 (blue), 2 (white), and 4 (red), and a possible incongruent answer is 3 (blue), 2 (white), and 1 (red). The pictures were taken with the camera of the HoloLens 2; the contrast of the digits appeared stronger to the participants. **Note:** The cube’s AR status information was presented within the participants’ peripheral vision, which required them to actively shift their gaze to process the information. The drawing is adapted with permission from [13].

#### 2.1.4 Data Preprocessing

The recorded data were preprocessed and analyzed offline using Matlab R2022a (The Mathworks Inc., Natick, USA), incorporating the open-source toolbox EEGLAB 2022.0 [24].

EEG from each participant was bandpass and bandstop filtered between 1 and 25 Hz and at 50 Hz, respectively (both Butterworth, 4th order, zero-phase). The downsampled signals (250 Hz) were segmented into epochs of 4 s, i.e., -1 to 3 s around the visualization onset (VIS), and corrected by subtracting the average amplitude of the baseline period [-0.6, -0.1] s. Next, we performed an independent component analysis by applying the extended infomax algorithm [25]. Components contaminated with artifacts, particularly ocular artifacts such as eye movements and blinks, were removed based on visual inspection [26]. To identify additional outliers in the EEG signals, we rejected epochs in which the amplitude of any channel exceeded ± 50 µV or had an abnormal joint probability or kurtosis, i.e., epochs for which either metric was greater than 5 SDs around the mean across epochs [27]. We also visually inspected the epochs. Furthermore, we computed the variance from each channel and spherically interpolated those exceeding the threshold, i.e., the third quartile plus 1.5 times the interquartile range. Finally, we extracted epochs relative to two time points of interest, i.e., (i) the onset of the AR visualization, i.e., the **VIS alignment**, and (ii) the first time point where the gaze intersected with the visualization, i.e., the **LOOK alignment**. After EEG data preprocessing, we had an average of 127.8 ± 8.4 congruent and 65.2 ± 3.9 incongruent epochs per participant and alignment left for further analyses. On average, we interpolated 1.7 ± 1.3 channels and removed 7.8 ± 2.9 independent components (all M ± SD). We used cluster-based permutation tests to identify significant group-level effects (*n* = 15) in the [-0.5, 1] s interval around both alignments [28]. For each channel-time point, we compared the EEG amplitudes with a two-sided t-test at a significance level of *α* = 5 %. Significant points were clustered based on temporal and spatial adjacency, and the cluster-level test statistic was computed as the sum of the t-values within each cluster. To assess statistical significance, we performed a cluster-based randomization procedure with 10,000 permutations, evaluated at a significance level of *α* = 5 %. This approach has been shown to effectively control the family-wise error rate when performing a large number of comparisons [29], as commonly necessary in EEG studies.

#### 2.1.5 Incongruity Decoding

The preprocessed epochs were used to classify the conditions, i.e., congruent versus incongruent, using shrinkage linear discriminant analysis (sLDA) [30]. Standard LDA classifiers trained on little data, as is common in ERP classification problems, often overestimate the eigenvalues of the real data distribution, leading to poor estimates of the covariance matrices and subsequently of the classifier weights. Shrinking covariance matrices has been shown to improve performance and has become a standard approach in BCIs for ERPs [31].

**Classifier Training and Testing** To train classifiers for each participant, we extracted time-domain features from the preprocessed EEG data within the [0.2, 1] s interval following the VIS and LOOK alignments, respectively. Prior research [32] demonstrated that averaging the amplitude of overlapping windows as features provides superior classification performance compared to other feature extraction approaches, e.g., using all samples within a window, in similar decoding tasks. We optimized the length of the overlapping windows for each participant using the data from all other participants. This procedure is described in *Hyperparameter Optimization*. Moreover, the default classification threshold of 0.5 was adjusted to account for the class imbalance in the data set and fixed to the prior probability of the minority class, as recommended by other works [33, 34]. Consequently, a trial was considered incongruent if its probability of belonging to the incongruent class exceeded 0.33.

Using these features, we iteratively trained and tested classifiers with the data of the participants not included in the hyperparameter optimization, following a stratified 10 × 5-fold (within-participant) cross-validation (CV) approach with random sampling. We computed the average true negative rate (TNR), true positive rate (TPR), and balanced accuracy (BA) on the 50 held-out test sets to summarize the decoding performance. The TNR is the ratio of correctly classified congruent trials to all congruent trials, and the TPR is its analog for incongruent trials. The BA is their average.

**Hyperparameter Optimization** We determined the optimal window length for each participant by evaluating five different durations, i.e., 0.1 to 0.5 s in increments of 0.1 s, with 50 % overlap. Specifically, we trained and tested sLDA classifiers using a stratified 5 × 5-fold CV scheme on the remaining 14 participants (leave-one-out), selecting the window that maximized the BA across all 350 folds (14 participants × 25 folds). We used the adjusted classification threshold of 0.33 for the optimization.

For the LOOK alignment, the 0.4 s window with 0.2 s overlap was selected for 14 participants, and once the 0.5 s window with 0.25 s overlap. The number of features per trial was 84 = 28 (channels) × 3 (windows: [0.2, 0.6], [0.4, 0.8], and [0.6, 1] s) for 0.4 s and 56 = 28 (channels) × 2 (windows: [0.2, 0.7] and [0.45, 0.95] s) for 0.5 s. For the VIS alignment, the 0.5 s window was selected for all participants following the same optimization procedure.

**Significance Level** For an infinite number of trials, the theoretical chance levels for TNR and TPR would equal the class proportions, approximately 67 % for the TNR and 33 % for the TPR. Given our limited data [35], we estimated the significance levels through a shuffling approach. For this purpose, we randomly permuted the class labels across trials 1,000 times and applied the same 5-fold CV scheme using LOOK-aligned data (sLDA with optimized window length and adjusted classification threshold). The individual significance levels were then defined as the 95th percentile of the test BA distributions (e.g., [36]).

### 2.2 Results

#### 2.2.1 Neurophysiological Results

The grand average neurophysiological responses, representing the mean ERPs of 15 participants, are summarized in Figure 3. For the VIS alignment (Figure 3b,c), no distinct brain activity can be observed during the baseline period, i.e., before visualization onset. There is a slight negative deflection in incongruent compared to congruent trials between 0.6 and 0.8 s. This parietal activity peaks after 0.77 s (-0.87 µV) but is not statistically significant (*p* = 0.15). After participants shifted their gaze toward the visualization (LOOK), the topographical distributions of the grand average ERPs show incongruity-related brain activity visible in the 0.28 to 0.42 s interval in centroparietal cortical areas (Figure 3e,f). In this segment, the difference signal incongruent minus congruent shows a negative response, i.e., an N400 effect, with a parietal peak after 0.36 s (-1.49 µV). Statistical analysis revealed significant activity in the corresponding channel-time point cluster (*p* = 0.02). The subsequent positive component of the difference signal, which peaks in Pz at 0.53 s (0.81 µV), fails to reach significance (*p* = 0.27). Condition-specific ERPs approximate 0 µV after 0.8 s.

#### 2.2.2 Classification Results

The binary classification results (congruent versus incongruent) for both alignments are summarized in Figure 4 and supplementary Tab. 1. When performing incongruity detection based on VIS-aligned data, we achieved an average TNR and TPR of 62.3 ± 4.7 % and 56.4 ± 6.1 %, respectively. For LOOK, both TNR and TPR increased to 72.8 ± 6.3 % and 67.3 ± 8.4 %. This results in BAs of 59.3 ± 4.9 % (VIS) and 70.0 ± 7.1 % (LOOK). The classification performance of LOOK is significantly better than that of VIS (average ΔBA = 10.7 %, *p <* 0.001, Wilcoxon signed-rank test). The average significance thresholds for the BA derived from the randomized classifier are 57.9 ± 0.4 % (all M ± SD). All participants performed better than chance with LOOK-aligned data, while only nine (60 %) could do so with VIS. This is indicated in Figure 4 (right) through blue or red (BA *>* significance level) and gray dots (BA ≤ significance level), respectively.

#### 2.2.3 Behavioral Results and Questionnaire

The distribution of the time it took participants to look at the visualization after the visualization onset is summarized in Figure 3d,g. The average time for congruent trials (LOOK C, 0.475 ± 0.09 s) did not significantly differ (*p* = 0.52) from the average time for incongruent ones (LOOK IC, 0.478 ± 0.08 s) (all M ± SD).

In a post-experimental questionnaire, participants were asked to report if they could identify incongruent counts (i) before looking at the visualization (i.e., while still looking at the cube) and (ii) when looking at the visualization. Hence, (i) refers to the time interval between VIS and LOOK, and (ii) refers to the time after LOOK. For both questions, participants could rate on a scale from 1 (“I could *always* identify incongruent counts”) to 6 (“I could *never* identify incongruent counts”). On average, participants reported being unable to identify incongruent counts pre-LOOK (5.4 ± 1.1) but could do so post-LOOK (1.1 ± 0.4) (all M ± SD). The paired difference between the two ratings is significant (Wilcoxon signed-rank test, *p <* 0.001).

**Figure 3.**
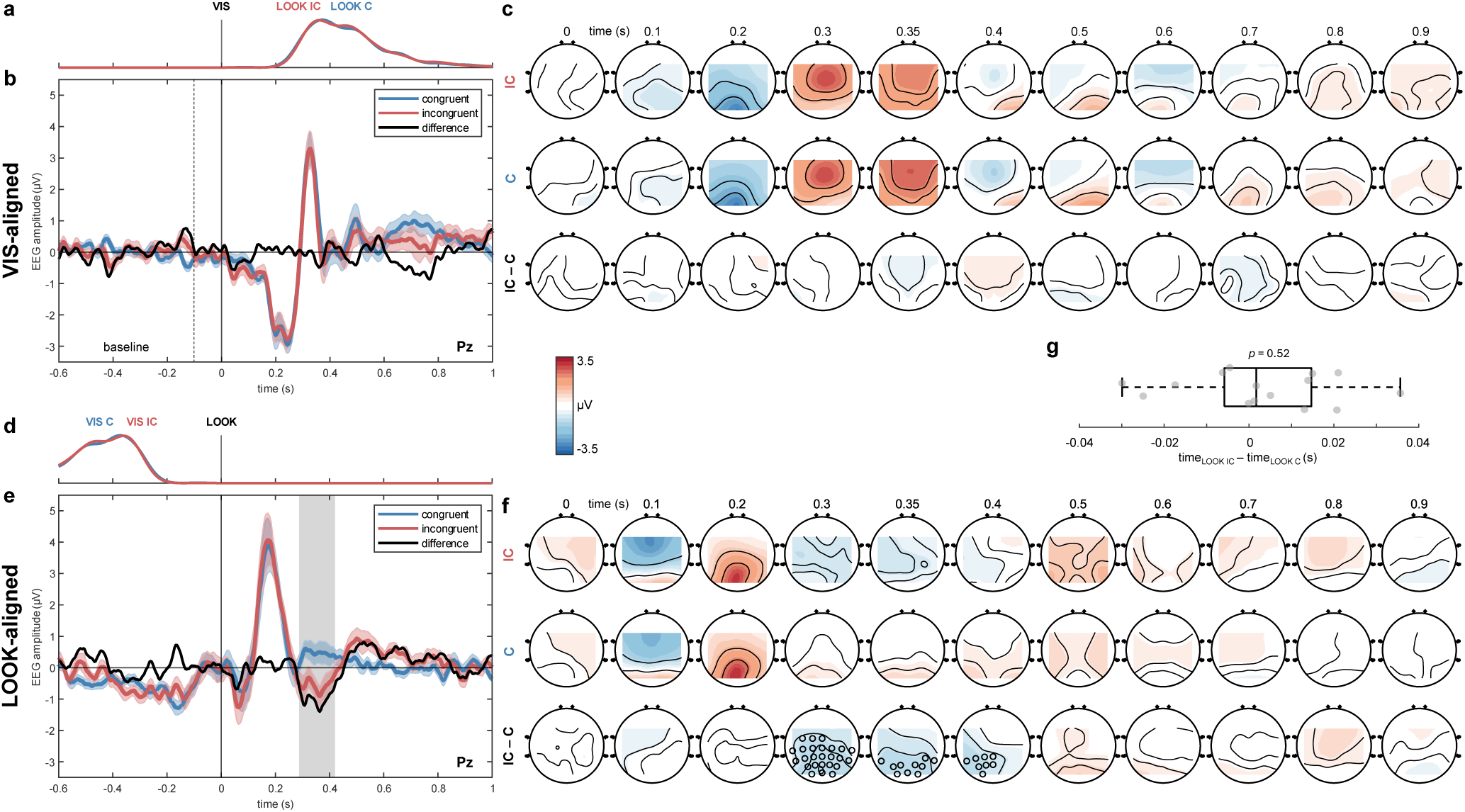
Condition-specific grand average EEG activity (*n* = 15) of VIS-(top) and LOOK-aligned (bottom) data for *Experiment 1*. **(a)** Distribution of the LOOK alignments relative to the visualization onset (VIS) at time = 0 s. The probability density of all trials from all participants was calculated using a Gaussian smoothing kernel with 0.03 bandwidth. **(b)** VIS-aligned grand average ERPs at channel Pz. Shaded areas indicate the standard error of the mean. The difference signal is computed as *incongruent (IC) minus congruent (C)*. No significant channel-time points were found (cluster-based permutation test, *p>* 0.05). **(c)** VIS-aligned grand average topographical maps at the indicated time points. No significant channel-time points were found. **(d)-(f)**As in **(a)-(c)**for the LOOK alignment. Significant channel-time points are indicated by the gray box in **(e)**and the circles in **(f)**, respectively (*p* = 0.02). **(g)** Boxplot summarizing the paired difference in the mean timings of the LOOK onsets across trials per participant. Condition-specific timings are not different between IC and C (*p* = 0.52, Wilcoxon signed-rank test).

### 2.3 Discussion

In **RQ1**, we investigated whether a combination of EEG and eye tracking data can be utilized to find brain activity related to semantic incongruity. We hypothesized that the low sampling frequency of the HMD’s eye-tracking system and the resulting uncertainty in the LOOK alignment [37, 38] would complicate the analysis of stimulus-related brain activity. However, the grand average EEG responses are consistent with prior research (e.g., [13]). We found significant clusters at approximately 0.3 to 0.4 s after LOOK, corresponding to the mismatch between the digital information and the cube, i.e., its physical context. Since participants could not identify incongruent counts before fixing on the visualization (as reported in questionnaires; see Section 2.2.3), no incongruity-related effect was observed in the VIS-aligned data. Consequently, classification with LOOK-aligned data significantly outperformed VIS-based decoders, achieving an average BA of 70 %.

Demonstrating the feasibility of eye tracking-informed EEG decoding is crucial for online incongruity detection. Commonly, online BCIs require continuous analysis and classification of ongoing brain activity (i.e., asynchronous classification), allowing users to engage in mental activities independent of external cues [18]. However, this strategy potentially inflates the number of false positive detections compared to synchronous classification approaches, where the evaluation of EEG signals is restricted to pre-defined time windows (e.g., following a marker) [20]. By incorporating gaze signals, we introduce a natural marker providing users with self-paced control of the system, while simultaneously constraining the EEG analysis to fixed windows.

**Figure 4.**
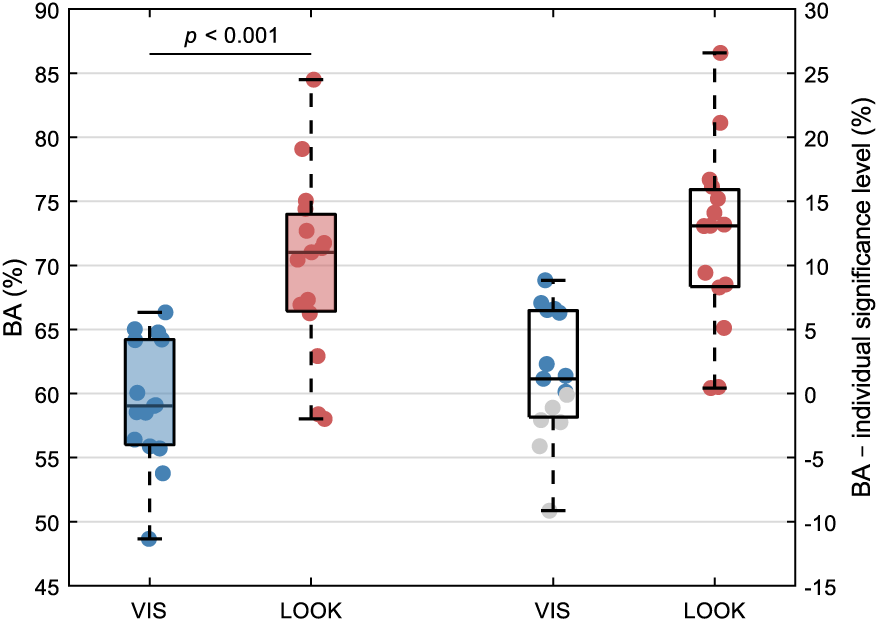
Participant-level classification results for *Experiment 1*. **Left:** BAs for VIS-(blue) and LOOK-aligned (red) data. BAs differ significantly (*p <* 0.001, Wilcoxon signed-rank test). **Right:** Differences between BAs and the individual significance levels (SLs). Blue and red dots indicate BA *>* SL, and grey dots BA *≤* SL.

## 3 Experiment 2

In *Experiment 2*, we addressed **RQ2** by extending the approach from *Experiment 1* to detect incongruent AR information online. Therefore, the experiment was divided into a calibration phase, in which personalized classifiers were trained, and a subsequent feedback phase, in which online incongruity decoding was performed.

### 3.1 Methods

#### 3.1.1 Participants

Fifteen healthy volunteers with normal or corrected-to-normal vision participated in the experiment (29.9 ± 3.1 years (M ± SD), 11 male, 4 female). All participants completed an online Ishihara pseudoisochromatic test to assess their color vision (as in Section 2.1.1). Five participants (33 %) reported having no prior AR experience, 6 reported being beginners or intermediates (40 %), and 4 reported being advanced or experts (27 %). Further, a total of 12 participants (80 %) had previously participated in at least one EEG study. The experiment was conducted in accordance with the Declaration of Helsinki (1975) and approved by the ethics committee of Graz University of Technology. All participants received written and oral instructions and signed an informed consent form. Participants received vouchers worth 20 euros as compensation.

#### 3.1.2 Experimental Setup and Procedure

The experimental setup was similar to *Experiment 1* and followed the description in Section 2.1.2. The platform on which the cube was positioned was extended with a panel to its right, allowing AR content to be presented on both sides of the cube, as shown in Figure 5b.

The experimental procedure lasted approximately 2 hours and consisted of three blocks. In the **first block**, a series of eye artifacts (blinks, horizontal and vertical saccades) and resting brain activity were recorded for about six minutes. A detailed description can be found in [39]. In the **second block**, i.e., the *calibration block*, participants performed 7 runs of 24 trials, 8 of which were incongruent and presented in a pseudo-randomized order (33 %). A detailed description of the trial structure can be found in Section 2.1.2. Contrary to *Experiment 1*, the AR visualization could either appear on the left or right side of the cube (in *Experiment 1*, it was always presented on the left). The location of the visualization was pseudo-randomized and equally distributed between both sides. The data of the calibration block were used to train the classifiers (see Section 3.1.4). The **third block**, i.e., the *feedback block*, consisted of 3 runs of 24 trials (8 incongruent, 33 %). In the feedback block, participants received online visual feedback based on the output of the classifier. Specifically, at the time point where the visualization disappeared in the calibration block, it was instead framed in green (classifier output is *congruent*) or red (classifier output is *incongruent*). The framed visualization disappeared after an additional 2 s (see Figure 5a). The remainder of the feedback trial structure is as in the calibration block.

Before the first block, we mounted the EEG electrodes and calibrated the HMD’s eye tracker, as explained in Section 2.1.3. Additionally, participants could familiarize themselves with the headset and the experimental paradigm and were informed about the occurrence and interpretation of the visual feedback.

**Figure 5.**
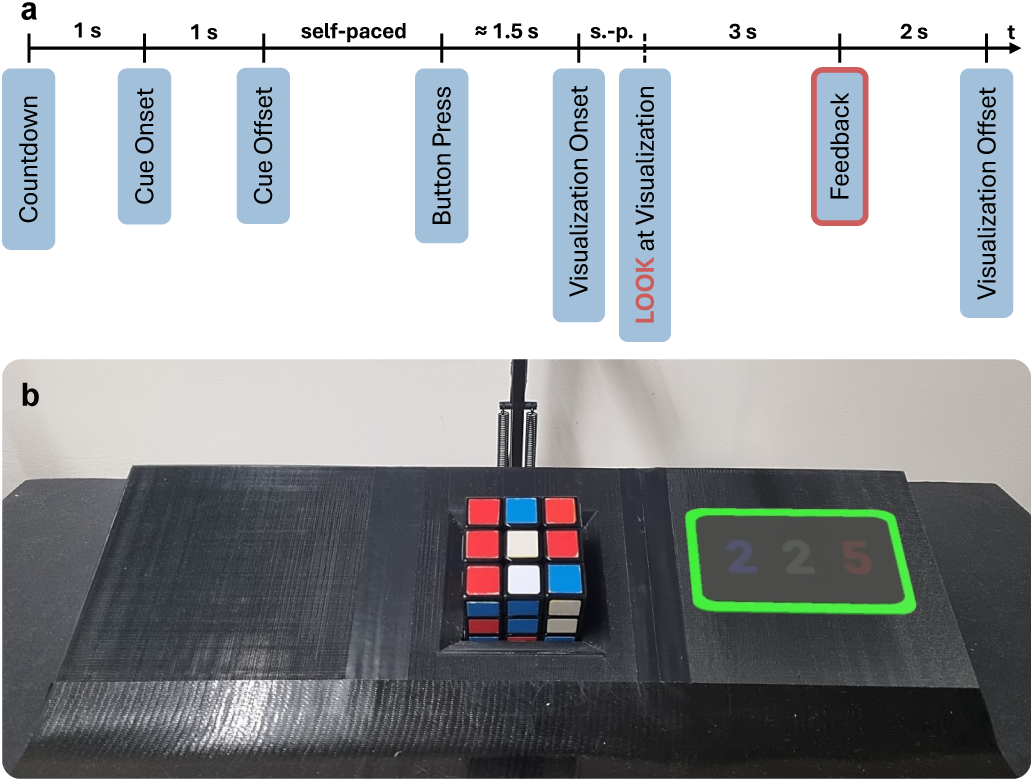
Experimental design of *Experiment 2*. **(a)** Timing of a single trial, including highlighted LOOK alignment and online feedback. Feedback was not given in the calibration block. **(b)** Adapted setup including AR information with visual feedback. The green frame indicates that the classification output of the current trial is of the class congruent.

#### 3.1.3 Data Acquisition and Preprocessing

The data acquisition is as in *Experiment 1* (see Section 2.1.3). The data preprocessing performed in the calibration and feedback blocks follows the same procedure. In each trial, the EEG data were cut into an epoch of 5 s, i.e., [-2, 3] s around the onset of the visualization, and downsampled to 250 Hz. Each trial was then bandpass and bandstop filtered between 1 and 25 Hz and at 50 Hz, respectively (both Butterworth, 4th order, zero-phase). Ocular artifacts, i.e., eye movements and blinks, were attenuated by applying the sparse generalized eye artifact subspace subtraction algorithm [39]. The algorithm was fitted using the data recorded in the first block. The preprocessed EEG signals of each trial were then further segmented into epochs of 2 s, i.e., [-0.5, 1.5] s with respect to the LOOK onset (as defined in Section 2.1.4). As in *Experiment 1*, we used cluster-based permutation tests (see Section 2.1.4) to identify significant group-level effects (*n* = 15) in the [-0.5, 1] s interval around LOOK, separately for the calibration and feedback blocks.

#### 3.1.4 Incongruity Decoding

**Classifier Training** After recording the calibration data in block 2, we trained a personalized sLDA classifier for the binary classification problem (congruent versus incongruent). For this, we used the preprocessed and LOOK-aligned calibration epochs. We rejected epochs with amplitudes exceeding ± 125 µV, with a kurtosis outside ± 3 SDs, or instances in which participants failed to look at the visualization within 1.5 s after its onset. On average, we rejected less than 1 % of the calibration epochs.

From the remaining epochs, we derived features from the time domain amplitude values within the [0.2, 1.25] s segment after LOOK. Specifically, we computed the averages of overlapping windows with window length *W* and overlap *W* /2. In a grid search scheme, we tuned the hyperparameters *W* ∈ {0.2, 0.25, 0.3, . . ., 0.7} s and classification threshold *T* ∈ {0.15, 0.175, 0.2, . . ., 0.5} to find the optimal settings for the online classification. For example, for *W* = 0.6 s, the classifier was trained with the average amplitudes of the windows [0.2, 0.8] s and [0.5, 1.1] s, resulting in 56 features (28 channels × 2 windows). In total, we evaluated 165 hyperparameter combinations (11 *W* × 15 *T*).

For each combination (*W*,*T*), we iteratively trained and tested classifiers using a stratified 2 × 5-fold CV approach with random sampling and computed the average TPR, TNR, and BA of the ten held-out test sets. We then smoothed the BA(*W*,*T*) using a 2D Gaussian smoothing kernel with SD = 0.5. The hyperparameter set that maximized the smoothed BA was chosen for the online classification. Consequently, we trained the final classifier with all calibration epochs that were not rejected using the optimized hyperparameters.

**Online Classification** We evaluated the classifier in the feedback block. The online data preprocessing followed the description in Section 3.1.3, and the classification features were derived based on the hyperparameters tuned in the *Classifier Training*. Participants received visual feedback based on the online classification output. If the probability of a trial belonging to the incongruent class exceeded the optimized threshold, the visualization was framed in red, otherwise in green. Subsequently, we computed the average BA of all feedback trials in which participants looked at the visualization within 1.5 s of its onset (99.4 % of the trials).

**Significance Level** Like in *Experiment 1*, we evaluated the classifier using a shuffling approach to compute personalized significance levels. The random classifier was trained on the calibration data using the optimized hyperparameters. We randomly permuted the class labels across calibration trials 1,000 times and tested each random classifier on the feedback trials. The individual significance levels were then defined as the 95th percentile of the test BA distributions.

### 3.2 Results

#### 3.2.1 Neurophysiological Results

The grand average ERPs of 15 participants are summarized in Figure 6. In the calibration data (Figure 6b,c), channel Pz exhibits two significant channel-time point clusters in the intervals [0.34, 0.46] s (*p* = 0.015) and [0.76, 0.84] s (*p* = 0.003). Within these clusters, the difference signal incongruent minus congruent shows a parietal minimum after 0.38 s (-1.53 µV), i.e., the N400. Additionally, we observed a late positive component with a central maximum after 0.79 s (1.51 µV).

In the feedback block, we found one significant channel-time point cluster in the segment between 0.27 and 0.48 s (*p* = 0.015). Within the cluster, the difference signal exhibits a parietal minimum after 0.46 s (-1.65 µV). Afterwards, a positive deflection in incongruent compared to congruent trials is visible but fails to reach significance (*p >* 0.9).

#### 3.2.2 Classification Results

The grand average classification results are illustrated in Figure 7. Figure 7a shows the results of the calibration block, including the average TNR (74.8 ± 6.6 %), TPR (74.8 ± 5.7 %), and BA (74.8 ± 4.8 %) achieved with the optimized hyperparameter sets (*W*,*T*), which are indicated with black dots. Further, the grand average results of each metric obtained with each hyperparameter combination are illustrated. The online classification performance is shown in Figure 7b. On average, a BA of 69.9 ± 5.3 % was achieved (left boxplot). The TNR and TPR are 72.0 ± 11.5 % and 67.8 ± 11.6 %, respectively. Fourteen out of 15 participants (93 %) reached accuracies above their individual significance levels (right boxplot, red dots), one did not (gray dot). The average significance level obtained with the random classifier is 62.5 ± 1.2 % (all M ± SD). The participant-level classification results are summarized in supplementary Tab. 2.

**Figure 6.**
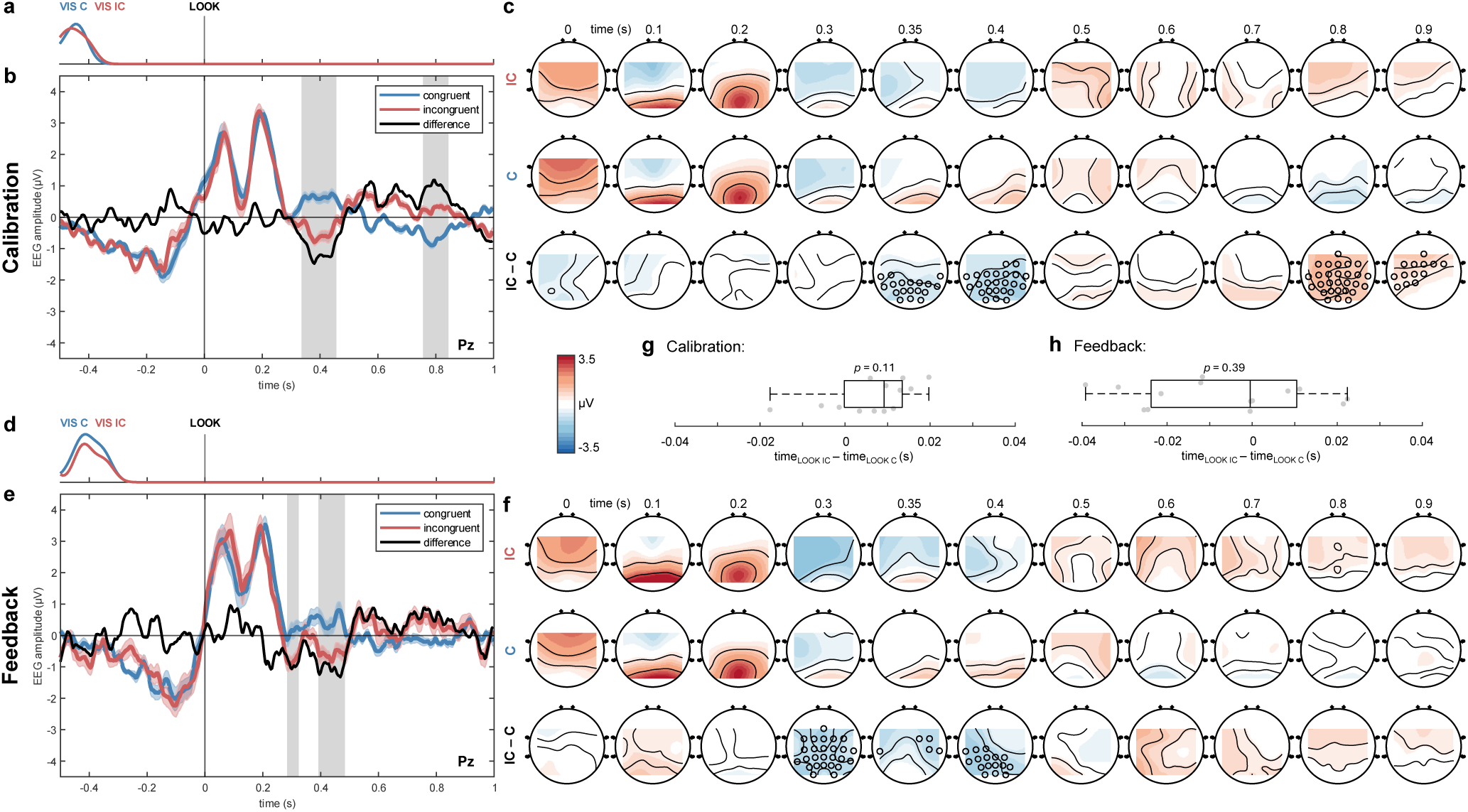
Condition-specific grand average EEG activity (*n* = 15) of the calibration (top) and feedback (bottom) blocks for *Experiment 2*. **(a)** Distribution of the visualization onsets (VIS) relative to the LOOK alignments (time = 0 s) in the calibration block. The probability density of all trials from all participants was calculated using a Gaussian smoothing kernel with 0.03 bandwidth. **(b)** Grand average ERPs of the calibration block at channel Pz. Shaded areas indicate the standard error of the mean. The difference signal is computed as *incongruent (IC) minus congruent (C)*. Significant channel-time points are indicated by the gray boxes (cluster-based permutation test, *p <* 0.05). **(c)** Grand average topographical maps of the calibration block at the indicated time points. Significant channel-time points are indicated (cluster-based permutation test, *p<* 0.05). **(d-f)** As in **(a-c)** for the feedback block. **(g)** Boxplot summarizing the paired difference in the mean timings of the LOOK onsets across trials per participant in the calibration block. Condition-specific timings do not differ significantly (*p >* 0.05, Wilcoxon signed-rank test). **(h)** As in **(g)** for the feedback block.

#### 3.2.3 Behavioral Results and Questionnaire

The time participants needed to shift their gaze from the cube to the visualization did not differ significantly between congruent and incongruent trials, either in the calibration (*p* = 0.11) or in the feedback block (*p* = 0.39). The grand average times for the calibration block are 0.449 ± 0.10 s and 0.453 ± 0.09 s for congruent and incongruent, respectively. The analogous grand averages for the feedback run are 0.430 ± 0.10 s and 0.432 ± 0.12 s (all M ± SD). The results are summarized in Figure 6g,h.

After the recordings, participants completed the same questionnaire as in *Experiment 1* (see Section 2.2.3). Again, the participants reported that they could identify incongruent counts (1.1 ± 0.3), but not before fixing on the visualization (5.5 ± 0.6) (all M ± SD). The paired difference between the two ratings is significant (Wilcoxon signed-rank test, *p <* 0.001).

**Figure 7.**
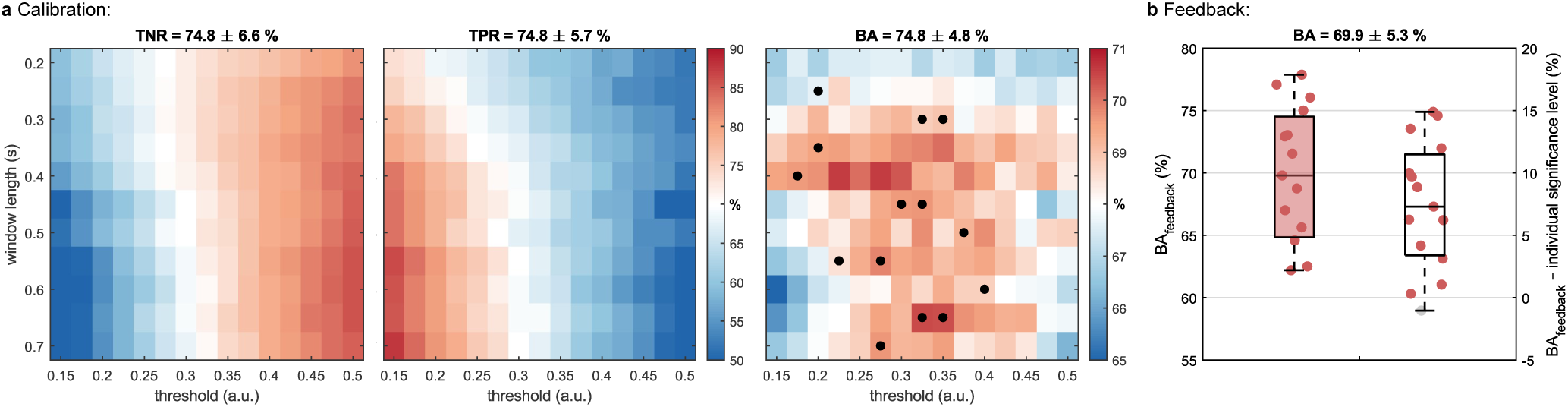
Classification results for *Experiment 2*. **(a)** Grand average TNR, TPR, and BA (*n* = 15) for each hyperparameter combination in the calibration block. Average metrics computed with the optimized hyperparameters are given (M ± SD). Black dots show the chosen hyperparameter pairs for each participant. **(b) Left:** Online classification results. Shown are the individual BAs in the feedback block. **Right:** Difference between the BAs and the individual significance levels (SLs). Red dots indicate BA *>* SL, and grey dots BA *≤* SL. The grand average BA is given (M ± SD).

### 3.3 Discussion

In **RQ2**, we investigated the feasibility of online incongruity detection by deploying the strategy developed in *Experiment 1*. We informed the BCI about the users’ visual attention through the HMD’s eye tracking system and decoded whether the corresponding AR information matched or mismatched their expectations. On average, the classification performance was approximately 70 %, matching the offline accuracies obtained in *Experiment 1*. Moreover, both experiments show consistent neurophysiological responses following the LOOK onsets. Minor variations can be attributed to differences in the paradigm (i.e., the visualizations could appear on either side) and the data processing pipeline, which had to be adapted for the online implementation. However, incongruity-related effects, specifically the N400 component, remained unaffected. Possible practical applications and limitations of online incongruity classification are summarized in Section 4.4.

## 4 General Discussion

### 4.1 Visualization Design

Panel designs, as utilized in the present study, are conventional visualizations semantically linked to the real scene. They are among the most frequently used situated visualizations in AR due to their flexibility and familiarity with many users [40]. When designing the AR visualization, we adhered to the principle of congruence [41] for compatibility between digital information and the real-world referent by relying on a common color code (i.e., blue, red, and white) and a consistent spatial structure (e.g., blue was always on the left). Additionally, the paradigm was designed such that participants were unable to differentiate incongruent counts before shifting their gaze toward them [42], reflecting real-world AR applications where complex information cannot be fully processed in peripheral vision [43]. In practical scenarios, AR often presents detailed and context-dependent data, such as intricate technical schematics [44], environmental overlays [45], or navigation cues [46], that require visual focus for accurate interpretation. As a result, users must shift their gaze to process data, making gaze-dependent information processing a crucial aspect of real-world AR interactions. In post-experimental self-assessments of both experiments, participants reported that they could not identify semantic mismatches before focusing on the data, confirming that the paradigm was suitable to simulate such scenarios.

### 4.2 Eye Tracking and Incongruity-Related Brain Activity

The ERP results of *Experiment 1* and *Experiment 2* are largely consistent concerning the N400 effect. Both show a negative deflection in incongruent trials, peaking in parietal channels after about 400 ms after LOOK (approximately -1.5 µV), while no incongruity-related responses are visible after VIS (as discussed in Section 2.3). A comparison to the results by Wimmer and colleagues [13], who investigated a very similar incongruity effect in a gaze-fixation task, reveals that the N400 effect observed in the present work is considerably weaker. Specifically, the difference in the reported peak amplitudes is approximately 1 µV. This discrepancy can be explained by differences in the presented stimuli. While the prior study featured incongruent counts with no clear connection to the cube’s current state (e.g., 1-6-1 or 9-9-9 instead of 4-2-3), we altered only one of the three digits (e.g., 4-5-3 instead of 4-2-3). In other words, the AR information presented in our study exhibits a higher degree of contextual relatedness. Indeed, relatedness has been shown to modulate the N400 in lexical priming tasks [47]. For example, “The pizza was too hot to *drink* ” elicits an N400 due to its unlikely, yet not entirely unrelated sentence completion (compared to “…*eat* ”), while the unrelated “…*cry* ” showed stronger amplitudes (e.g., no associative, semantic, categorical, or phonological relationship). Consequently, the obvious relation between the incongruent counts and the cubes’ state led to the observed reduction of the N400 amplitude. The amplitude difference may be partially attributed to a smearing effect in the grand average ERPs caused by the jittered LOOK alignments, as discussed in Section 2.3 [37, 38]. However, the spatial distribution of the N400, peaking in parietal areas of the brain, remained.

Participants consistently interpreted incongruent counts as semantic mismatches within their physical context. However, this observation is not trivial. Prior research has demonstrated that erroneous human-machine interactions elicit ErrPs, i.e., stimulus-related brain activity following erroneous events, which considerably differ in amplitude, latency, and topography. It has been shown that ErrPs are elicited not only when making a mistake but also by machine errors, such as misinterpreting user input [48, 49]. ErrPs are usually characterized by a frontocentral error-related negativity followed by a centroparietal error positivity within the first 400 ms after the error event [50, 51]. Given these findings, one could hypothesize that participants would perceive incongruent counts as machine errors resulting from erroneous color detection. However, the observed latencies and scalp distributions of the elicited ERPs indicate that semantic mismatch, rather than machine error, was the dominant effect in the present work.

### 4.3 Eye Tracking-Informed BCIs Can Decode AR Incongruity Online

Since participants could not identify incongruities before gazing at the augmentation, we hypothesized that classifiers trained with VIS-aligned data would not perform above chance. Indeed, the average BA (59.3 %) is only marginally better than random classification (57.9 %). Conversely, decoders trained with LOOK-aligned trials performed significantly better and achieved a mean TNR of 72.8 % and TPR of 67.3 %, yielding a BA of 70.0 % on average. This is an improvement over previous works: there are only a handful of papers that have demonstrated N400 single-trial classification toward BCIs [52], many of them in linguistic paradigms that typically present a sequence of prime words followed by matching or mismatching probe words. Reported offline accuracies are in the range of approximately 60 % [53, 54, 55] to 65 % [13] correctly classified instances. However, a direct comparison is difficult as considerable differences in paradigm design, type of incongruity, number of trials, data preprocessing, and classification methodology exist.

Regarding the latter aspect, prior research could not identify considerable differences between the decoding performance of traditional classification algorithms, e.g., sLDA or support vector machines, and neural networks, such as EEGNet [55, 56]. This could be a consequence of the limited amount of data typically acquired in EEG experiments, which restricts the potential performance advantages of deep learning approaches [57]. However, the classification performance achieved in this work is within the range of more extensively studied effects in practical BCIs, such as error processing [50]. With our classification strategy, we could transfer the offline accuracies of *Experiment 1* to an online scenario (*Experiment 2*). The visual feedback provided in the online block is a proxy for any system response that might occur after detection. Notably, the presented strategy relies on eye tracking signals that can be acquired with most commercial HMDs [58], thereby avoiding additional system complexity.

### 4.4 Applications and Limitations

This work provides substantial steps toward BCIs for real-world AR applications by enabling online decoding of incongruity while granting users autonomous control over when data are presented and allowing free eye movement across the scene. This approach does not aim to assess the objective validity of information but rather the user’s perception of it, which opens the door for manifold applications. For instance, in industrial settings, monitoring sensors for unusual values is crucial for detecting system anomalies. While N400 decoding is not typically used for such monitoring, unless interpreting the values requires expert knowledge, it could be valuable to assess if experts and novices perceive and interpret these values differently in training scenarios. If perceived incongruities are identified, the system could adapt autonomously by presenting the data in an alternative format to mitigate misinterpretations or by providing additional context when the information appears unintuitive. Additionally, the system could flag incongruent augmentations, enhancing the system design to improve reliability and user experience. More broadly, this approach can be transferred to numerous scenarios that require the processing or interpretation of AR visualizations, ranging from simple real-world annotations to learning tools in education.

Nevertheless, we acknowledge that the proposed approach still presents challenges that should be addressed in future research. First, the current approach is limited to a fixed data representation (Arabic numerals), a single physical referent (a Rubik’s Cube), and a constant complexity (three digits, with one differing by three). Two recent studies combining EEG and AR demonstrated that similar effects can be observed across different data visualizations (e.g., bar charts [13]) and various target objects [15]. However, it remains unclear how the neural responses – and consequently the decoding performance – are affected in scenarios that require interpreting more complex data or when incongruities become more subtle. Additionally, distracting environments may impact the perception of AR data, potentially altering the associated brain activity. Therefore, EEG responses should be examined across diverse scenarios to identify the most promising conditions for system adaptation based on physiological reactions.

Beyond usability, classification performance is a major consideration for practical BCIs. In this work, we could show that incongruity-decoding is possible with accuracies similar to other commonly considered ERPs in BCIs, e.g., approximately 70 to 80 % for error decoding [50]. These results are promising, given that the N400 effect is particularly affected by high inter-participant variabilities and low detection rates within individual participants [52, 59]. Nonetheless, BCIs need to decode brain activity more accurately to be relevant in practice. Various approaches exist to mitigate this issue; they range from hybrid BCI approaches that fuse information from multiple physiological or non-physiological sources [60, 61] to data augmentation [62] and transfer learning [63] to inflate the amount of training data. Particularly, transfer learning is a compelling approach as it allows classifiers to be trained on data from other users or tasks, thereby reducing the time required for user-specific training before BCI applications [64].

## 5 Conclusion

In this work, we successfully demonstrated, for the first time, an online N400-BCI for AR applications. The development of this online BCI encompassed two experiments, in which we implemented an interactive paradigm simulating a common AR application, i.e., the presentation of in-situ information related to a physical referent. In the first experiment, we showed that eye-tracking markers can support the detection of incongruent AR information, such as information that contradicts the user’s expectation. This approach extends prior literature on brain responses to augmentations by allowing users to exhibit natural visual behavior during AR interactions. This strategy was applied in a second experiment, where eye-tracking guided BCIs decoded EEG signals online. In both experiments, we observed consistent neurophysiological responses, specifically the N400 effect, following the visual fixation of incongruent information. Subsequently, we could decode this effect with an average online classification accuracy of approximately 70 %.

The proposed approach raises the possibility of informing systems whether AR information appears counterintuitive or unexpected to users, regardless of the objective validity. This allows for autonomous system adaptation, which could be utilized to proactively provide additional context to make the information more accessible to users or aid system diagnosis by flagging unexpected encounters. Future efforts should be directed towards (i) investigations across different data representations, information complexities, and degrees of incongruity to assess their influence on the physiological responses, (ii) improving the accuracy of incongruity decoders, and (iii) their application in out-of-the-lab scenarios.

## Acknowledgments

The Know Center is funded within the Austrian COMET Program - Competence Centers for Excellent Technologies - under the auspices of the Austrian Federal Ministry of Innovation, Mobility and Infrastructure, the Federal Ministry of Economy, Energy and Tourism, and by the State of Styria. COMET is managed by the Austrian Research Promotion Agency FFG. This work was supported by the TU Graz Open Access Publishing Fund.

## Author contributions

**Michael Wimmer:** Conceptualization, Methodology, Investigation, Formal Analysis, Writing - Original Draft, Writing - Review & Editing, Visualization. **Neven ElSayed:** Conceptualization, Writing - Review & Editing. **Bruce H. Thomas:** Conceptualization, Writing - Review & Editing. **Gernot. R. Müller-Putz:** Conceptualization, Writing - Review & Editing, Supervision. **Eduardo E. Veas:** Conceptualization, Writing - Review & Editing, Supervision, Funding Acquisition.

## Supplementary data

Supplementary tables and video are provided.

## Footnotes

1 https://www.colorblindnesstest.org/ishihara-test/

2 https://github.com/kkoomen/qbr

