## Supplemental Tables for "An Online Brain-Computer Interface for Detecting Incongruity in Augmented Reality Applications"

**Table 1:** Participant-level classification results for *Experiment 1*. For the VIS- and LOOK-alignments, TPR, TNR, and BA, as well as the significance level (SL), are given.

| Participant | **VIS** | | | **LOOK** | | | SL (%) |
|---|---|---|---|---|---|---|---|
| | TNR (%) | TPR (%) | BA (%) | TNR (%) | TPR (%) | BA (%) | |
| E1-P01 | 58.7 | 52.8 | 55.7 | 62.7 | 54.1 | 58.4 | 58.0 |
| E1-P02 | 53.3 | 44.0 | 48.7 | 66.4 | 59.4 | 62.9 | 57.8 |
| E1-P03 | 61.8 | 58.3 | 60.1 | 69.3 | 63.2 | 66.3 | 57.8 |
| E1-P04 | 65.5 | 63.0 | 64.2 | 83.8 | 85.2 | 84.5 | 57.9 |
| E1-P05 | 63.7 | 54.3 | 59.0 | 80.8 | 69.3 | 75.0 | 58.9 |
| E1-P06 | 62.1 | 55.0 | 58.5 | 72.2 | 61.6 | 66.9 | 58.6 |
| E1-P07 | 57.5 | 50.0 | 53.8 | 69.1 | 65.5 | 67.3 | 57.9 |
| E1-P08 | 58.9 | 53.9 | 56.4 | 63.5 | 52.6 | 58.0 | 57.5 |
| E1-P09 | 68.4 | 61.6 | 65.0 | 82.0 | 76.2 | 79.1 | 58.0 |
| E1-P10 | 70.7 | 62.0 | 66.3 | 73.7 | 71.7 | 72.7 | 57.5 |
| E1-P11 | 62.4 | 55.8 | 59.1 | 77.3 | 71.5 | 74.4 | 57.7 |
| E1-P12 | 65.9 | 51.1 | 58.5 | 74.0 | 66.9 | 70.4 | 57.4 |
| E1-P13 | 57.1 | 54.6 | 55.9 | 71.8 | 70.2 | 71.0 | 58.0 |
| E1-P14 | 67.1 | 62.5 | 64.8 | 74.2 | 68.5 | 71.4 | 58.2 |
| E1-P15 | 60.5 | 67.8 | 64.2 | 70.7 | 72.8 | 71.4 | 57.6 |
| M ± SD | 62.3 ± 4.7 | 56.4 ± 6.1 | 59.3 ± 4.9 | 72.8 ± 6.3 | 67.2 ± 8.4 | 70.0 ± 7.1 | 57.9 ± 0.4 |

**Table 2:** Participant-level classification results for *Experiment 2*. For the calibration block, the TNR, TPR, and BA achieved with the optimized hyperparameters $W$ and $T$ are shown. For the feedback block, the online TPR, TNR, and BA, as well as the significance level (SL), are given.

| Participant | **Calibration** | | | | | **Feedback** | | | |
|---|---|---|---|---|---|---|---|---|---|
| | TNR (%) | TPR (%) | BA (%) | W (s) | T (a.u.) | TNR (%) | TPR (%) | BA (%) | SL (%) |
| E2-P01 | 71.5 | 79.3 | 75.4 | 0.25 | 0.2 | 70.8 | 58.3 | 64.6 | 63.5 |
| E2-P02 | 74.5 | 71.5 | 73.0 | 0.65 | 0.35 | 64.6 | 75.0 | 69.8 | 60.9 |
| E2-P03 | 67.4 | 82.2 | 74.8 | 0.40 | 0.175 | 64.6 | 87.5 | 76.0 | 62.5 |
| E2-P04 | 67.9 | 72.1 | 70.0 | 0.55 | 0.275 | 64.6 | 66.7 | 65.7 | 61.5 |
| E2-P05 | 85.3 | 83.0 | 84.2 | 0.30 | 0.325 | 70.2 | 54.2 | 62.2 | 61.9 |
| E2-P06 | 73.3 | 82.2 | 77.8 | 0.35 | 0.2 | 75.0 | 62.5 | 68.8 | 62.5 |
| E2-P07 | 81.8 | 68.7 | 75.3 | 0.60 | 0.4 | 58.3 | 66.7 | 62.5 | 63.5 |
| E2-P08 | 75.2 | 73.1 | 74.2 | 0.30 | 0.325 | 50.0 | 84.0 | 67.0 | 60.8 |
| E2-P09 | 78.5 | 74.1 | 76.3 | 0.45 | 0.325 | 95.7 | 60.0 | 77.9 | 63.0 |
| E2-P10 | 77.9 | 77.7 | 77.8 | 0.30 | 0.35 | 75.0 | 75.0 | 75.0 | 63.0 |
| E2-P11 | 66.8 | 70.5 | 68.6 | 0.65 | 0.325 | 68.1 | 75.0 | 71.5 | 61.9 |
| E2-P12 | 78.1 | 81.3 | 79.7 | 0.55 | 0.225 | 83.3 | 70.8 | 77.1 | 62.5 |
| E2-P13 | 61.9 | 65.7 | 63.8 | 0.70 | 0.275 | 73.3 | 72.7 | 73.0 | 63.0 |
| E2-P14 | 81.3 | 72.3 | 76.8 | 0.45 | 0.3 | 87.5 | 41.7 | 64.6 | 61.5 |
| E2-P15 | 80.5 | 68.2 | 74.3 | 0.50 | 0.375 | 79.2 | 66.7 | 72.9 | 65.6 |
| M ± SD | 74.8 ± 6.6 | 74.8 ± 5.7 | 74.8 ± 4.8 | 0.47 ± 0.1 | 0.30 ± 0.1 | 72.0 ± 11.5 | 67.8 ± 11.6 | 69.9 ± 5.3 | 62.5 ± 1.2 |
